# Functional and historical drivers of leaf shape evolution in Palms (Arecaceae)

**DOI:** 10.1101/2021.10.26.465896

**Authors:** Maria Fernanda Torres Jimenez, Nicolas Chazot, Thaise Emilio, Johan Uddling Fredin, Alexandre Antonelli, Soren Faurby, Christine D Bacon

## Abstract

**Aim:** Leaves display a remarkable variety of shapes, each with potential ecological advantages in specific climates. While relations between leaf shape and either climate or height has been relatively well studied in eudicots, the macroecological drivers of shape remain poorly known in monocots. Here, we investigated associations between climate and plant height with the evolution of leaf shape in a clade with high species and morphological diversity.

**Location:** Global.

**Time period:** Cretaceous to contemporary.

**Major taxa studied:** Palms (Arecaceae).

**Methods:** We apply a Bayesian phylogenetic mixed model to test for associations between climate and leaf shape (all entire-leaved, pinnate-dissected, palmate-dissected, and costapalmate). We further reconstruct the ancestral leaf shape using multistate speciation and extinction models and compare the frequency of shapes with global temperatures through time.

**Results:** We find that plant height associates with dissected leaves and that annual precipitation associates with pinnate shapes. The ancestral leaf shape is unclear but early diversification was dominated by pinnate-dissected palms, which has remained the most species-rich form of leaves throughout palm history.

**Main conclusions:** Palms that are tall and live in humid regions are more likely to have pinnate leaves. Through geological time scales, temperature did not play an obvious role in determining leaf shapes. This study contributes to our understanding of how the diversity of leaf shapes is linked to biological and climatic factors.

## 1. Introduction

Leaves are the engines of most life on land. They capture atmospheric carbon dioxide and convert it into accessible nutrients that sustain ecosystem functioning and health. The basic functions they carry out – photosynthesis, transpiration, and respiration – depend on temperature and water availability. Because they carry out such critical functions, leaves are likely under strong natural selection, resulting in morphological adaptations to environmental conditions (Nicotra et al., 2011). Throughout angiosperm evolution, variation not only in length and width but also in blade dissection, has evolved numerous times independently (Nicotra et al., 2011). Nonetheless, the drivers of leaf shape evolution and its adaptive potential are poorly known, with research being limited to a handful of species or to model eudicots (Nicotra et al., 2011; Chitwood & Sinha, 2016; Leigh et al., 2017), hindering generalisations at a larger taxonomic scale (Conklin et al., 2019). The evolution of leaf shape results from trade-offs between physiological and allometric constraints, phylogenetic history, and natural selection (Givnish, 1987; Tsukaya, 2006; Nicotra et al., 2011). Therefore, understanding the extent to which leaf shapes have evolved in response to climate, allometry, or natural selection, and how they have changed over time and across geographies, may shed further light on the evolution of life forms and biological adaptations.

General expectations regarding leaf function can be drawn from examining their adaptations in different environments. Traits like leaf dissection, length and effective width (diameter of the largest circle drawn within a leaf margin; Leigh et al., 2017) vary across climates, which affects plant temperature regulation and interacts with water availability (Nicotra et al., 2011; Peppe et al., 2011). In environments with extremely hot temperatures, plants tend to have small or dissected leaves to avoid reaching damaging temperatures (Leigh et al., 2017; Wright et al., 2017). Deeply dissected leaves effectively function as a collection of small leaf units, with faster heat loss through convection than entire-leaved ones of the same area (Givnish, 1979; Gurevitch & Schuepp, 1990), and are less likely to reach damaging temperatures when exposed to extreme heat. Thus, shapes that reduce damage at high temperatures are expected to be beneficial in hot and dry environments (Nicotra et al., 2007; 2008; Leigh et al., 2017). Under drought conditions where water for transpiration is limited, stomata close and compounding high temperatures threaten leaf function. In such cases, smaller or deeply dissected shapes reduce overheating and optimise leaf safety (Nardini & Luglio, 2014). Species with large and wide leaves, optimised for high gas exchange, could benefit from dissection if water transportation to blade areas farthest from the rachis is more effective (Givnish, 1979). In contrast, cool climatic conditions may favour entire-leaved over dissected, where leaf heating is not extreme and photosynthetic surfaces can be wide.

Leaf shape and size may also be optimised to maximise light capture. This is particularly relevant for monopodial plants (non-branching plants with a single growth axis) or understory species that depend on leaf size, angle, or number to increase the total photosynthetic area (Givnish, 1988; Valladares et al., 2002; Renninger & Phillips, 2016). However, leaf size is limited by allometry (the proportional change in size of different parts of an organism). Corner’s rule states that “the larger and sturdy an axis is, the larger and more complicated are its appendages” (Corner, 1949; Tomlinson, 2006). Dissecting large leaves enables independent movement of leaflets and reduces wind drag (Niklas, 1992; Vogel, 2009). Similarly, the “rapid growth hypothesis” links dissection and allometry (Givnish 1984; Niinemets, 1998), and states that dissection (or compound leaves) enables fast growth in high light by investing in longer rachises, which is energetically cheaper than wood density and branching (Malhado et al., 2010). To maximize light capture in the absence of branching, elongated petioles or rachises can be used as alternative strategies (Givnish, 1988), facilitating optimal trade-offs between leaflets’ angle and self-shade (Valladares et al., 2002). When petiole lenghts are similar across species, elongated rachises could be effective for outstretching leaves or leaflets to radii farther from the plant stem. In addition, elongating light structures like the rachis could solve limitations imposed by supporting heavier structures. In the absence of branching, alternative strategies for maximising light capture could be elongated petioles or rachises (Givnish, 1988); particularly rachises for outstretching leaves or leaflets to radii farther from the plant stem and facilitating optimal trade-offs between leaflets’ angle and self-shade (Valladares et al., 2002). Moreover, long petioles for better light capture, elongating light rachises (and thus the leaf blade) can solve limitations imposed by supporting long and heavy petioles.

Here, we use palms (Arecaceae) to understand the macroevolutionary drivers of leaf shape variation. Palms are an ideal group to address this question given their wide distribution and morphological diversity. They are tropical and subtropical, with 80% of the species distributed within a 15-30°C mean annual temperature range (Dransfield et al., 2008), and exhibit wide leaf shape variation (pinnate, palmate, and costapalmate, all of which can be entire-leaved or dissected; Dransfield et al., 2008; Horn et al., 2009). Palms are primarily monopodial with non-deciduous leaves (Tomlinson, 2006; Dransfield et al., 2008), allowing us to control for the effect of branching strategies over leaf-variable relationships. Understanding the evolution of leaf shape through past and current environmental conditions provides a context for predicting plant responses to changing climates (Chitwood & Sinha., 2016).

In this study, we aim at disentangling the contributions of climatic (extrinsic) and allometric (intrinsic) factors on palm leaf shape by testing four hypotheses: **H1:** Climate, and in particular temperature, contributes more to leaf shape than allometry. Dissected species (pinnate- and palmate-leaved) should be found at higher temperatures (or lower precipitation and higher aridity) than entire-leaved species regardless of height. **H2:** Allometry, likely plant height, contributes more to leaf shape. We test H1–H2 by comparing i) extant dissected and non-dissected shaped species and ii) all shapes in a pairwise manner. **H3:** Either plant’s height in relation to forest canopy height or elongated rachises contribute to leaf shape by trading light capture maximisation and leaf support, particularly when comparing pinnate and palmate species. Longer rachises (in pinnate species) translate into long blades and more surface for light capture in cases for which petioles and stems are short. We test H3 by comparing all shapes in a pairwise manner. **H4:** If climate is a strong contributor to leaf shape evolution through time, this should be reflected in a temporal congruence between leaf shape shifts and major global climatic changes since the Late Cretaceous, when palms are estimated to have originated (Couvreur et al., 2011; Baker & Couvreur, 2013). We test these hypotheses by estimating variable effects and estimating ancestral character states.

## 2. Methods

We conducted Generalised Linear Mixed Models (GLMMs) and ancestral trait estimation analyses at the species level using a time-calibrated Maximum Clade Credibility tree generated from the tree distribution in Faurby et al. (2016) and updated by Hill et al. (2021). The phylogeny included 2550 species for which we annotated leaf shapes, recovered geographic records from the Global Biodiversity Information Facility (GBIF), and estimated species medians for the climatic and allometric variables. We standardised the taxonomy across all data sources using Kew’s World Checklist of Selected Plant Families (WCSP) for Arecaceae (Govaerts et al., 2020) and removed records that could not be unambiguously assigned to accepted species.

### Leaf shape in palms

Leaf shape variation in palms can be described by three features: size, plication and dissection. Since plication is phylogenetically conserved, we focused on dissection and shape, but see supplementary material for a brief description of size and plication. Leaves are either dissected or entire-leaved. Dissected leaves can be pinnate, palmate, or costapalmate depending on rachis length and the presence of a costa (here treated as the equivalent of rachis in pinnate leaves but it is an extension of the leaf axis; Dransfield et al., 2008). Polymorphism, where intraspecific variation in leaf shape occurs, only involves either pinnate-entire or pinnate-dissected shapes. Genomic analyses (e.g. Loiseau et al., 2019) confirm most intraspecific variation as true polymorphisms within populations, rather than separate taxa grouped under one species name.

We classified species by dissection (dissected vs. entire) and shape (pinnate, palmate, and costapalmate) based on Genera Palmarum II (Dransfield et al., 2008), additional information on the PalmWeb (https://palmweb.org/, last consulted January 2023), and herbarium specimens (access through GBIF). We merged the costapalmate and palmate shapes because the only difference between them is the presence of a costa. We removed the bipinnate category from the GLMM analyses but merged them to pinnate for ancestral state estimation because bipinnate includes only 14 out of 2550 species. Of the species analysed, the majority are pinnate (66.18 %; Table S1, Supplementary Material), followed by palmate+costapalmate (21.26 %) and entire (5.12%). Only 75 (6.76 %) species are polymorphic and while they were included in the ancestral state estimation, they were excluded from the GLMMs because they cannot be assigned to a unique shape category (Table S2).

### Palm allometry data

We used Palm_Traits v.1 (Kissling et al., 2019) to calculate plant height by adding the variables ‘MaxStemHeight_m’, ‘Max_Petiole_length_m’, and ‘Max_Blade_Length_m’, with ‘MaxStemHeight_m’ set to zero for acaulescent species. We estimated an index that measures plant height controlled by canopy height (height over canopy) as a proxy to wind-drag exposure and the Understorey/Canopy variable in Kissling et al. (2019). The index differentiates between e.g. two tall species, one occurring in a high canopy and another in a low canopy forest. Values of height over canopy are higher than 2.70e+10 for tall species in low canopies, close to one for species with the same height as the canopy, and smaller than 2.65e-08 for short species under high canopies (based on the 0.25 and 0.75 quantiles). To calculate it, we divided the palm height species value by the height extracted at the coordinate point from the Global Vegetation Height layer (Simmard, 2011). Calculations and code are available at doi.org/10.6084/m9.figshare.21230453. We annotated 61-100% of the species in the phylogeny. Species without annotations had no data in Palm_Traits v.1, were climbing, or had both climbing and no-climbing habits (479 species, Table S3); climbing species were removed from the GLMM analyses because their life strategies differ from other palms and their stem height is not comparable.

### Climatic data

We downloaded 994,084 raw occurrences from GBIF (last consulted in April 2022) and excluded fossils and records without coordinates. We kept observations, living and preserved specimens, and material samples. We used the R package (R Core Team, 2018) ‘CoordinateCleaner’ v2.0-20 (Zizka et al., 2019) to remove duplicate coordinates per species and records nearby science institutions, bodies of water, and city/country centroids, using a buffer of 5000 m and 10000 m for centroids and cites respectively. We used a Python script (doi.org/10.6084/m9.figshare.21230453) and the World Geographical Scheme for Recording Plant Distributions maps (TDWG; Brummitt, 2001) to remove records falling outside their native botanical countries according to the WCSP. We obtained 124,703 clean records representing 61.56% of the 2550 species in the phylogeny (Table S4), which we used to extract all Chelsa v2.1 bioclim variables (Karger et al., 2017; Karger at al., 2018) with a python script (doi.org/10.6084/m9.figshare.21230453). Finally, we estimated the species medians for every variable from which we could extract the information (Table S4).

We determined correlations between variables and chose those with a Spearman’s coefficient −0.7 < LJ < 0.7 (Fig. S1). We additionally estimated the variance inflation factor for all variables using python’s ‘statsmodels’ v0.13.5 (Seabold and Perktold, 2010) and kept those with values below two. We kept five climatic and three allometric variables: mean annual temperature (°C*100), temperature seasonality (standard deviation °C*100), annual precipitation (kg m-2); precipitation seasonality (standard deviation kg m-2), aridity index (mean annual precipitation/mean annual potential evapotranspiration; kg m-2/time unit), plant height (m), rachis length (m), and height over canopy. Temperature seasonality and mean annual precipitation, and plant height and height over canopy were correlated (Spearman’s correlation coefficient = −0.8 and 0.73 respectively; Fig. S1); however, we assessed them in separate models to determine if the effect of e.g. temperature is larger than e.g. precipitation on leaf shape. All variables used were log_10_-transformed and standardised to have zero mean and a standard deviation of one (Table S4).

### Generalised linear mixed models

We fit a series of GLMMs for the first three hypotheses using logistic regressions and pairwise comparisons; entire (0) vs. dissected (1), pinnate (0) vs. palmate (1), entire (0) vs. palmate (1), and entire (0) vs. pinnate (1); these were implemented in the R package ‘MCMCglmmRAM’ v2.24 (Hadfield, 2015). The GLMMs assess shape rather than shape evolution, thus, we treat pinnate-entire and palmate-entire as entire-leaved, pinnate-dissected as pinnate, and palmate-dissected as palmate. For computational constraints, we ran every model on 42 (Adams, 1979) phylogenies randomly selected from the distribution, each as a random effect. For every model, we ran four independent chains of 10,000,000 iterations, a thinning of 5,000, and 8000 burn-in. Analyses were run on a HPE Cray OS computer through the Swedish National Infrastructure for Computing. We estimated effective sampling sizes (ESS) and evaluated chain convergence with a python script (doi.org/10.6084/m9.figshare.21230453). Chains converged and ESS scored higher than 200. We evaluated the significance of the effects for every predictor based on whether the 2.5 and 97.5% quartiles of the estimated density overlapped zero.

### Ancestral state estimation

To reconstruct the evolution of leaf shapes we used the Multi-State Speciation and Extinction (MuSSE) model from the R package ‘diversitree’ v0.9-16 (FitzJohn, 2012). We adjusted the leaf shape dataset to avoid over parametrization of our models: 1) excluded plication due to high phylogenetic clustering (Fig. 1; but see Couturier et al., 2011); 2) used a “polymorphic” category for species exhibiting more than one shape; 3) excluded all species with missing data for leaf shape. The final dataset included 2543 species. Ancestral state estimation concerns shape and dissection evolution, thus we designed three different model structures – two binary state structure excluding polymorphic, pinnate vs. palmate, entire vs. dissected, and a five-state structure (pinnate-entire, pinnate-dissected, palmate-entire, palmate-dissected and polymorphic).

**Figure 1.**
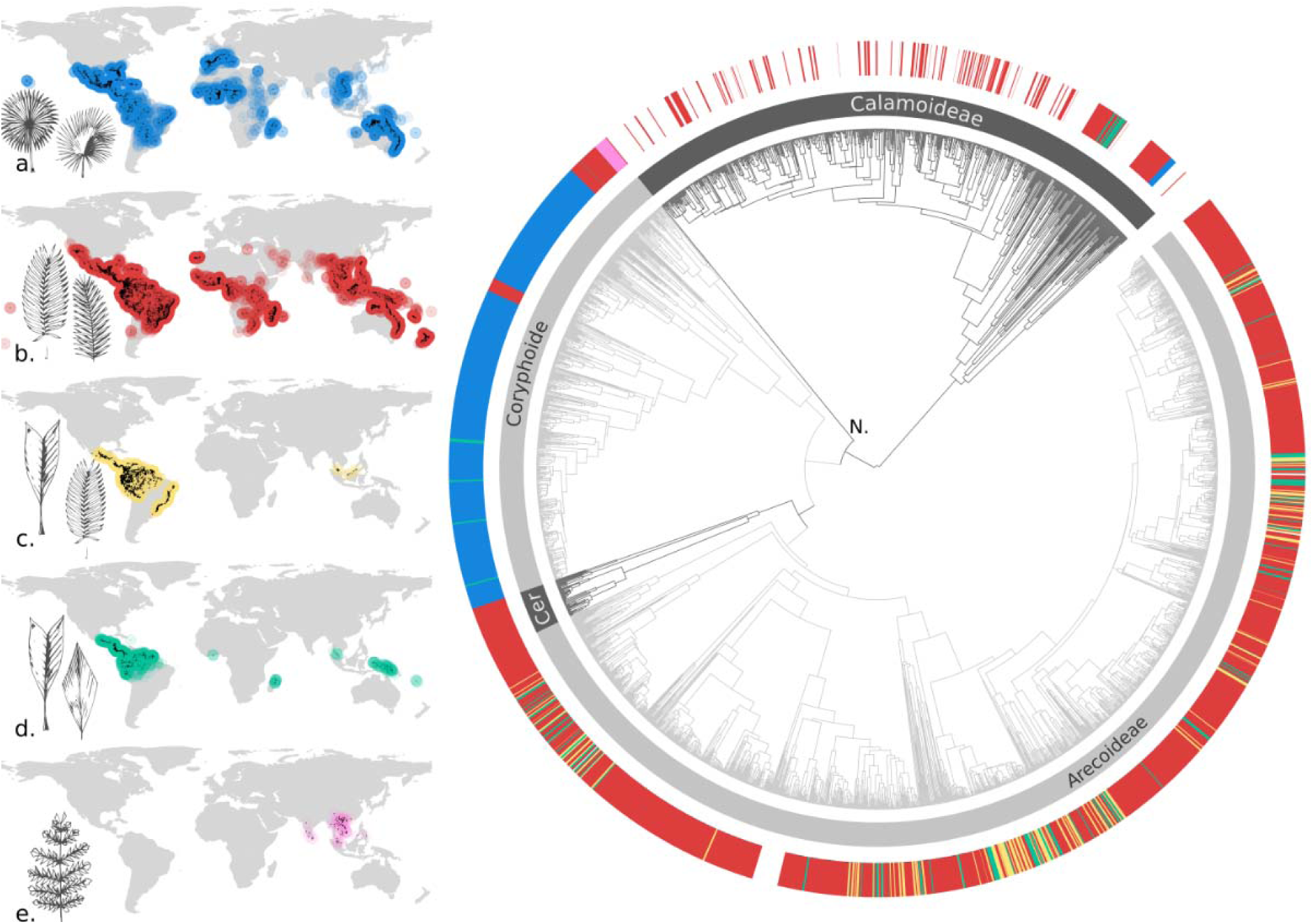
Leaf shape distribution at a global scale and across the Palm phylogeny. **Left:** The maps show the approximate distribution of shapes based on cleaned GBIF records (see Methods). Leaf silhouettes, originally by Marion Ruff Sheahan, were modified from Genera Palmarum (Dransfield et al., 2008). **Right:** The calibrated maximum clade credibility tree of 2550 palm species (Faurby et al., 2016; Hill et al., 2021). Dark and light greys are used to distinguish between subfamilies. The outermost ring shows the distribution of leaf shapes across the phylogeny and follows the same colour scheme in the maps. Species with no leaf shape colours are climbing and were removed from the Generalised Linear Mixed Models. **a)** Palmate + Costapalmate, **b)** Pinnate, **c)** Polymorphic, **d)** Entire, **e)** Bipinnate. **Cer.=** Ceroxyloideae. **N.=**Nypa.

For the two binary structures we fitted all possible parameter constraints using maximum likelihood (ML) parameter estimations and compared them from AIC scores. For the five-state structure, we refined the basic model structure by assuming constraints on transition rates. We did not allow simultaneous change of shape and dissection (*e.g.*, from palmate-entire to pinnate-dissected). As polymorphism only includes pinnate leaves, we assumed direct transitions between palmate and polymorphic states to be impossible and constrained the corresponding transition parameters to zero. Given the large number of parameters remaining (22), we selected the best-fitting model using a backward model selection procedure using ML estimation on the maximum clade credibility tree only. Once we obtained the best-fitting model for the two binary and the five-state structures, we used it to perform an MCMC analysis to estimate the posterior distribution of parameter estimates. We ran the MCMC for 3,000,000 generations, sampling every 3,000 generations and a burnin of 10%. We used the ML parameter estimation as starting points for the MCMC. In addition, we performed ML estimations of the best-fitting model (1) for 100 trees randomly sampled from the posterior distribution to account for phylogenetic uncertainty and (2) for a range of starting parameters. More details about model selection and parameter constraints are available in the Supplementary Material.

For all model structures, we used the best-fitting MuSSE model to estimate ancestral states, and the relative frequency of leaf shape through time by sampling the relative probabilities of shape estimated for each node 100 times. For each iteration, at each branch where a state transition occurred, we sampled a random timing from a uniform distribution for the event along that branch. Finally, for each iteration, we counted the number of transitions within a 5 million-year sliding window and calculated the rate of transitions through time by dividing the number of transitions by the sum of branch lengths within each time interval. We repeated this procedure for 100 trees randomly sampled from the posterior distribution. Finally, we compared the relative shape frequency through time with the global temperature change scale computed for an ice-free ocean (Zachos et al. 2001; Condamine et al., 2020).

## 3. Results

### Leaf shape and variable distributions

Pinnate and palmate species tend to be larger than entire-leaved species with median plant heights of 7.39 m and 7 m, respectively, while entire-leaved species had a median height of 3.8 m (Table S5), leaf and petiole lengths included in all cases. Dissected shapes together had a median height twice that of non-dissected species. Pinnate-leaved species had a median rachis length of 2.9 m and a median height over canopy of 4e+10, while palmate-leaved species had a median rachis length of 2.4 m and a median height over canopy of 7e+10. Entire-leaved species had a median height over canopy of 1.3e+10. Pinnate and palmate species tend to be equal or taller than the canopy, whereas entire-leaved species were rarely tall enough to approach the canopy and do not occur in open habitats. alms are widely distributed across climatic gradients except for habitats where soil temperatures fall below −2LJ for a long period of time. Pinnate and palmate species are distributed farther from the equator with wider climatic seasonality, while the distribution of entire-leaved species is concentrated in tropical areas near the equator where annual temperature and precipitation are more constant. Pinnate and palmate species have wider median distributions for most climatic variables than entire-leaved species, except for narrower annual mean temperature and temperature seasonality ranges in palmates. The median of the distribution for mean annual temperatures is highest for palmate species, then entire-leaved species, and lowest for pinnate species (Table S5, Supplementary Material).

### Generalised Linear Mixed Models

We found that plant height was positively associated with dissected leaves, particularly pinnate shapes. The association is significant when comparing entire-leaved vs. pinnate-leaved species (P; proportion of samples above/below zero=0.99; Fig. 2, Table S8) and entire-leaved vs. dissected species (P=0.99). The pattern was consistent across models regardless of which climatic variables were included. For the model including annual precipitation instead of annual temperature seasonality, annual precipitation showed a positive and almost significant association with pinnate species when compared to entire-leaved species (P=0.952), but not when comparing dissected and entire-leaved shapes (P=0.89). Nevertheless, the effect of plant height is an order of magnitude larger than the effect of precipitation (1.58E+05 and 6.68E+04 respectively, Table S8, Supplementary Material). For models comparing pinnate and entire-leaved shapes in which height over canopy was included instead of plant height (these variables are correlated and were not included in the same model), only height over canopy was almost significantly and positively associated with pinnate shapes (P=0.953). Height over canopy had a similar effect as plant height on leaf shape for pinnnate vs. entire-leaved models (1.02E+05 and 1.58E+05). However, the significance of height over canopy’ effect was lost if the model did not account for annual precipitation. No climatic variables were consistently and significantly associated with leaf shape (0.35<P< 0.94) and all variables were not significant in models involving palmate shapes (Fig. 2, Table S8).

**Figure 2.**
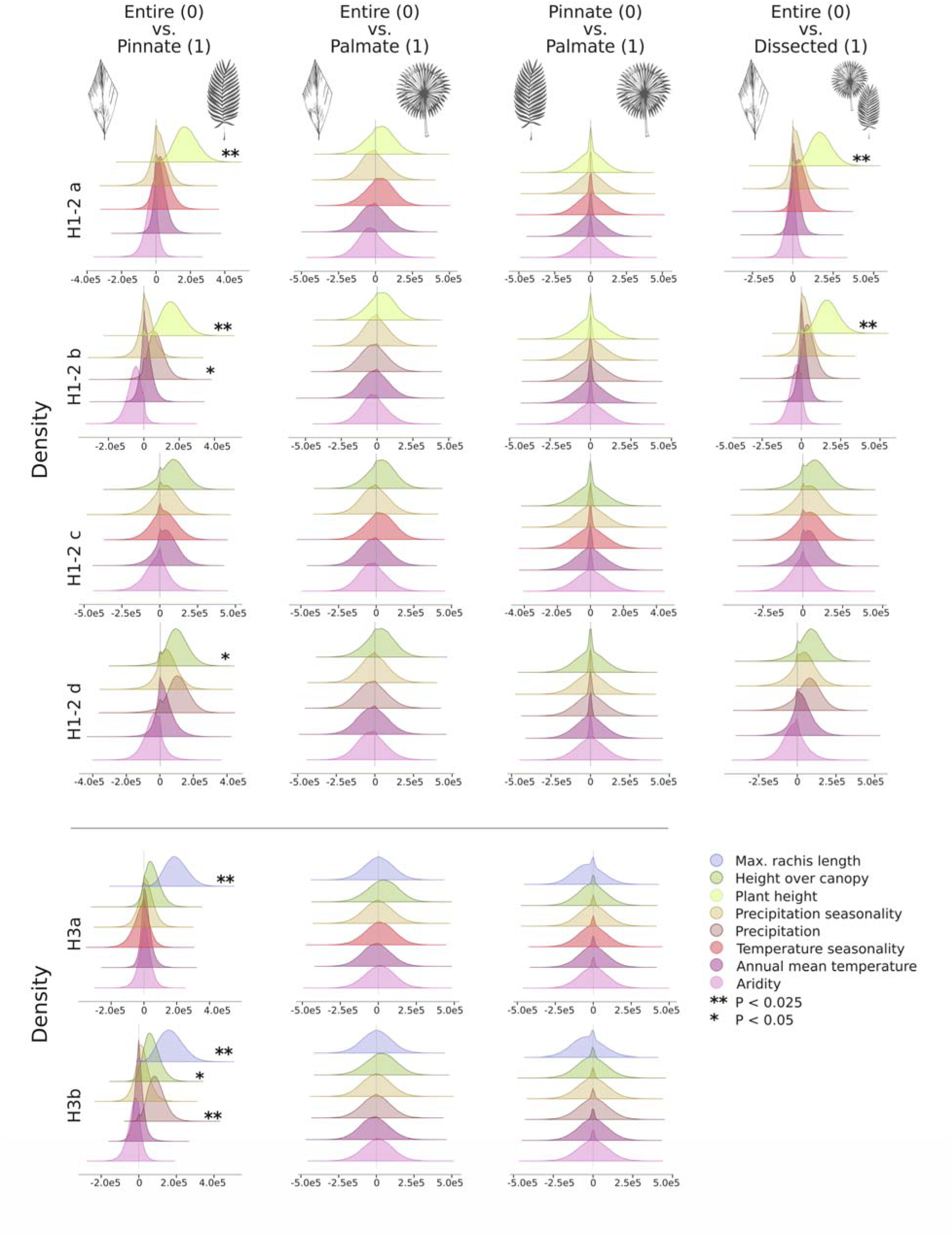
Estimate distributions from the Generalised Linear Mixed Models (GLMMs). Leaf shapes show the pair compared in the models (logistic regression; left=0, right=1). Each row shows the results from the models concerning hypotheses H1-3. Palmate = (costapalmate+palmate). We tested hypotheses H1-2 for each shape separately and for entire vs. dissected (palmate + costapalmate + pinnate), H1-2a, H1-2b, H1-2c, and H1-2d correspond to the models including either plant height or height over canopy, or temperature seasonality or precipitation. Hypothesis H3 was tested for each shape separately and H3a and H3b correspond to the models including either temperature seasonality or precipitation. Leaf silhouettes, originally by Marion Ruff Sheahan, were modified from Genera Palmarum (Dransfield et al., 2008).

When disentangling the effects of rachis length, height over canopy, and climate, we found a significant positive association between rachis length and pinnate shapes when compared to entire-leaved species; plants with longer rachises are more likely to be pinnate. In addition to rachis length, annual precipitation had a significantly positive association with pinnate species and height over canopy was almost significant (P=0.99 and P=0.968, respectively). Here, the effect of rachis length was one order of magnitude larger than precipitation and height over canopy, which had similar effects. The models resulted in no association between rachis length and shape when comparing pinnate and palmate species (0.34< P<0.5; Fig. 2, Table S8).

### Ancestral state estimation

The best-fitting model for dissection vs. entire-leaved state estimation had six parameters (Fig. S3). The speciation rate (lineage/unit of time, l^-1^my^-1^) for entire-leaved lineages was more than twice higher as that of dissected lineages and the extinction rate was simultaneously much lower for entire-leaved lineages, but transition rates were highly asymmetrical favouring transitions from entire-leaved to dissected than the reverse (Fig. S8). For pinnate vs. palmate model structure, we found no difference in speciation or extinction rates between shapes. Transitions from pinnate to palmate leaves were three times higher than the reverse transition, but both were very low (Fig. S9). The best-fitting model for the five-state model had 12 parameters (Fig. S5). Most notably, the speciation rate of polymorphic species was more than twice higher as that of the others. The extinction rate for the polymorphic and palmate-dissected lineages was much lower than that of pinnate-dissected and palmate-entire lineages. The resulting net diversification was highest for polymorphic lineages, followed by pinnate-entire leaves. Both palmate-entire and pinnate-dissected net diversification rates were close to zero. Transitions were highest from polymorphic leaves to pinnate-dissected, twice higher than transitions from pinnate-entire to polymorphic leaves and near 10 times higher than transitions from polymorphic to pinnate-entire leaves. All other types of transition rates were very low (Fig. S10). For all three model structures, we obtained identical parameter estimations from varying starting parameters, a posterior distribution of trees or MCMC analyses.

Across a posterior distribution of 100 trees, the two-state structures estimated the root to be pinnate and entire-leaved (Fig. S6-7). According to the five-state structure, the shape at the root was most likely polymorphic or pinnate-entire (Fig. 3). However, we found the results to poorly represent the pattern of early diversification of shapes. The relative frequency of shapes through time, calculated from the ancestral state estimations, shows instead that pinnate-dissected leaves dominated through the history of palms, representing between 50 to 80% of all lineages. Despite appearing as likely leaf shape at the root, entire-leaved lineages diversified only during the second half of the Cenozoic. According to our estimations, palmate-dissected palms appeared about 75-85 My ago and represented about 20% of palm lineages throughout history. Polymorphic lineages were poorly represented during the middle Cenozoic, despite being estimated as a probable shape for the root of the tree and increasing in frequency during the last 30 My (Fig. 4).

**Figure 3.**
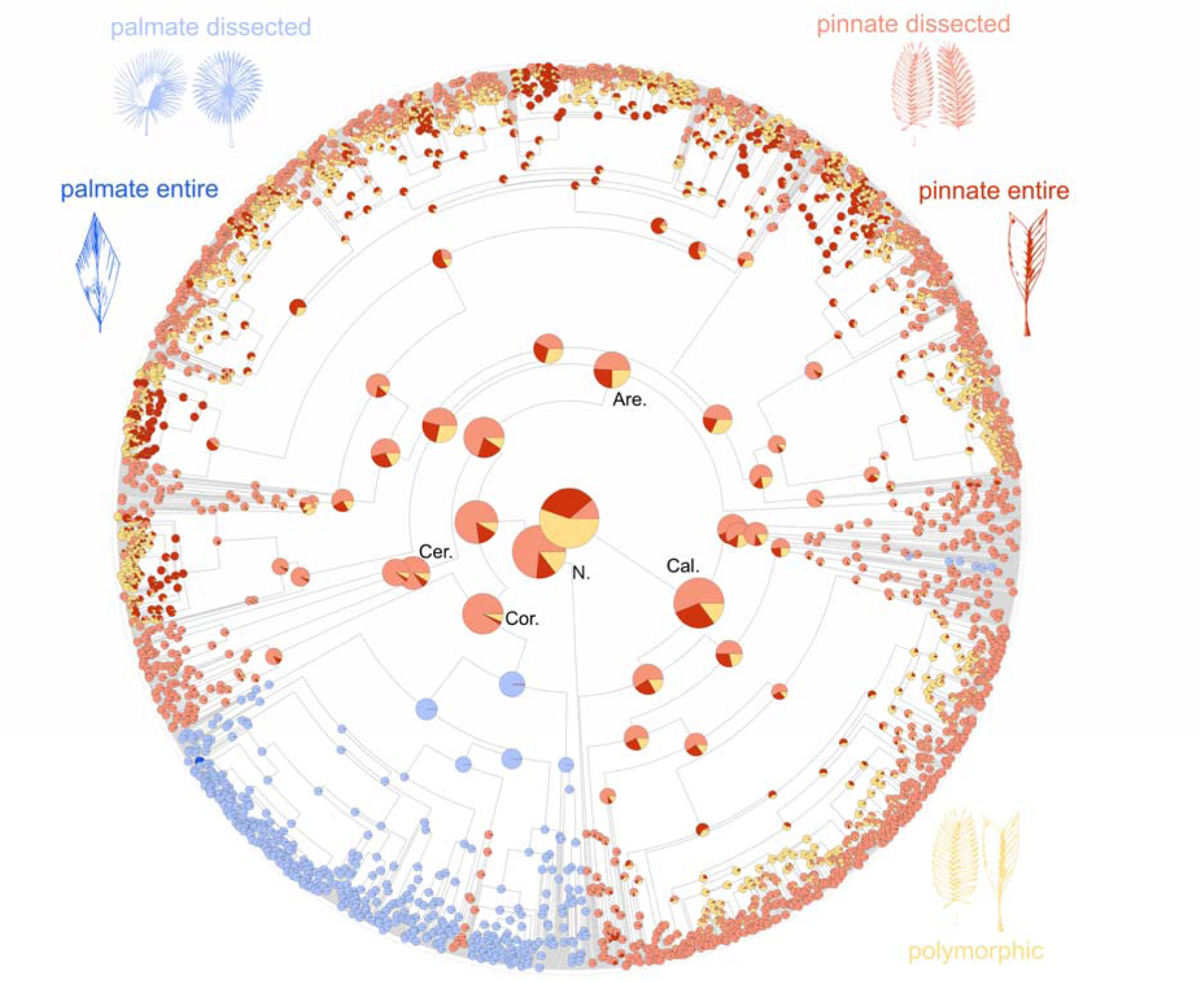
Ancestral state estimation of leaf shape across palms (Arecaceae) using the calibrated maximum clade credibility tree (Faurby et al., 2016; Hill, et al., 2021). The pie charts show the probability of each state at the given node. States are colour-coded as follows: Blue: Palmate (costapalmate+palmate); Red: Pinnate; Green: Entire; Yellow: Polymorphic (where individuals of the same species have either entire or pinnate leaves). Leaf silhouettes, originally by Marion Ruff Sheahan, were modified from Genera Palmarum (Dransfield et al., 2008). **N.=**Nypa, **Cal.**=Calamoideae, **Cor.**=Coryphoideae, **Cer.**=Ceroxyloideae, **Are.**=Arecoideae.

**Figure 4.**
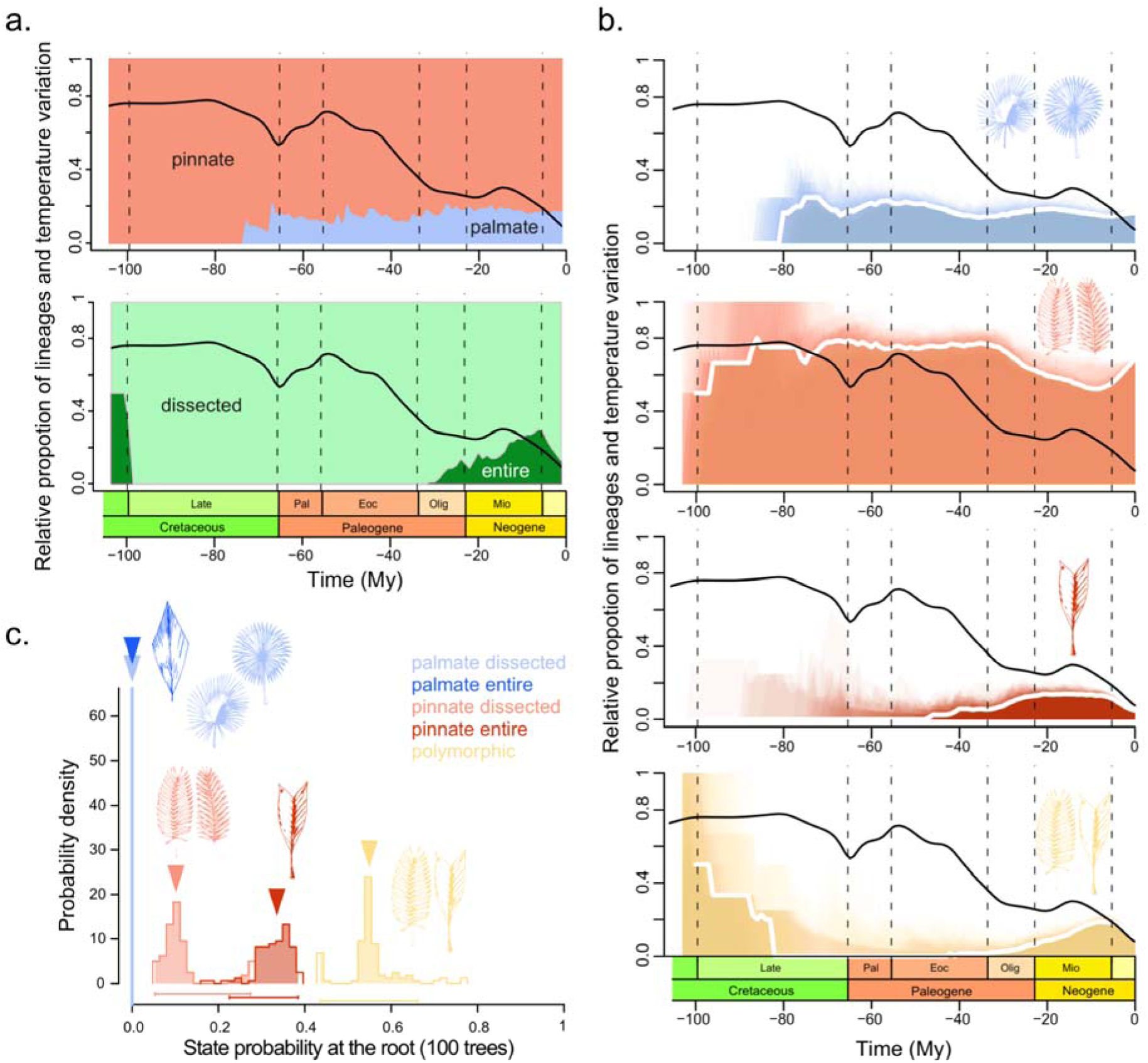
Palm leaf shape through time. **(a). Top:** Relative proportion of lineages with either entire-leaved or dissected shapes compared to global temperature variation through time. **Bottom:** Relative proportion of lineages with either pinnate or palmate (palmate+costapalmate) shapes compared to global temperature variation through time. **(b).** Relative proportion of lineages with (from top to bottom) palmate-dissected leaves, pinnate-dissected leaves, pinnate-entire leaves, or are polymorphic (intraspecific variation between pinnate-entire and pinnate-dissected). The proportion of palmate-entire lineages is close to zero and not plotted. In **(a)** and **(b)**, the white lines show the average proportion and the coloured lines show the relative proportion sampled from the 100 ancestral state estimations. **(c).** Relative probability of the ancestral state at the root of Arecaceae across 100 random trees sampled from the posterior distribution of the MCMC based on the five-state model. The horizontal bars at the bottom show the width of the posterior distribution and triangles show the default starting parameter. Leaf silhouettes, originally by Marion Ruff Sheahan, were modified from Genera Palmarum (Dransfield et al., 2008).

## 4. Discussion

In this study, we aimed at disentangling the contributions of plant allometry and climate on leaf shape by testing whether **H1)** temperature contributes more than allometry to leaf shape; or **H2)** allometry, as plant height, contributes to shape, or **H3)** rachis length or height over canopy contribute to shape; and whether **H4)** there is temporal congruence between shape frequency and major global temperature changes. This suggest that taller plants are more likely to have dissected leaves and, regarding climate, only annual precipitation contributes and makes species more likely to have pinnate leaves (Fig. 2). We also found that longer rachises make pinnate leaves more likely. In congruence with our results, we observed a lack of correspondence between leaf shape frequency and temperature fluctuations through time. Our models of ancestral trait estimation revealed that the ancestor of all palms likely had entire-leaved and pinnate or polymorphic leaves (Fig. 3–4c).

### Leaf dissection and climate

Contrary to H1, neither mean annual temperature nor temperature seasonality had an effect on leaf shape (Fig. 2, Table S8). We expected that plants exposed to extremely high temperatures would have dissected leaves as a strategy to avoid overheating their leaves faster (Nicotra et al., 2007; 2008; Leigh et al., 2017). Similarly, we expected that a high aridity index – a proxy for a lack of water in the environment– would favour pinnate leaves over other shapes. Our results showed no significant associations between climate and leaf shape for most of the models and only the effect of annual precipitation was significant when comparing pinnate and entire-leaved species, thus rejecting **H1**. Species that experience high annual precipitation are more likely to be pinnate than entire-leaved when plant height is also present in the model. Since plant height and annual precipitation are not correlated, our reuslts suggests that even though the effect of plant height is greater, annual precipitation and plant height both seem to explain independent variation.

Moreover, the association between climate and palmate shapes is not significant (Fig. 2); three non-mutually exclusive hypotheses could explain the general lack of associations, of which the first two are more likely. First, the discrete shape categories in our models might not fully capture interspecific variation in dissection depth in palmate species and could be insufficient for unveiling the climate-palmate associations. Second, leaf traits unrelated to shape could be key adaptations to different environments (Horn et al., 2009). For example, the loss of non-lignified fibre bundles in the leaf mesophyll mostly observed in coryphoid palms or the presence of bridge-like veins connecting the adaxial and abaxial layers of the leaf are traits thought to be advantageous in dry environments (Kenzo et al., 2007; Horn et al., 2009). Third, in the case of palmate and costapalmate species, the correlation between climate and shape is difficult to disentangle from a phylogenetic effect due to the strong geographic structuring of related lineages, particularly in the Coryphoideae subfamily to which the majority of palmate and costapalmate species belong.

### Leaf shape and plant allometry

We expected plant height to influence leaf shape via allometry so that large plants would have large and more likely dissected leaves to avoid mechanical damage (Corner, 1949; Chazdon, 1991). We found that plant height has a positive effect on dissection and taller plants are more likely to have pinnate leaves (Fig. 2, Table S8). Our results were consistent when comparing dissected –pinnate and palmate together– and pinnate species against entire-leaved species, supporting **H2**. Only when comparing palmate- vs. entire-leaved species, all effect estimates were not significant (Fig. 2). This suggests that the association between plant height and dissection is primarily driven by the number of pinnate species (66.18% vs. 21.26% species with palmate leaves) and that other factors are involved when palmate species are considered (as discussed in the section above).

Following the “rapid growth hypothesis” (Givnish, 1984; Niinemets, 1998), we expected plant height to relate to dissection. Malhado et al. (2010) tested this hypothesis in Amazonian species (except palms and polymorphic eudicots), and found an association between dissection and both low wood density and rapid diameter growth. They argue that dissection is adaptive under favourable light conditions because producing compound or dissected leaves is physiologically less expensive since enlongater rachises are cheaper than supporting structures (petioles in the case of palms) and thus promotes rapid vertical growth (Malhado et al., 2010). We found a positive association between height over canopy and pinnate shapes when these are compared to entire-leaved shapes (Fig. 2, Table S8), although marginally significant. The significance of this association, however, should be interpreted carefully as height over canopy values are derived from, and correlated with, plant height. The lack of association between height over canopy when comparing pinnate and palmate-leaved species is explained by the large proportion of species occupying similar spaces in the canopy where overall light availability conditions are similar. Malhado et al. (2010) excluded palms from their analyses and testing whether rapid growth or light capture optimisation better explains dissection is not possible with our data; a deep look at species-level measurements of growth rates and light capture efficiency could shed a light on this question. Nonetheless, our results suggest that allometry is more important than light capture in palms and further exploration of this association is worthwhile in future studies.

Finally, we expected light capture optimisation to drive rachis length and contribute to shape when comparing pinnate vs. palmate species. Pinnate leaves differ from palmate leaves in that pinnae attach to a long axis and not a centroid. On one hand, the elongated rachis could be a strategy for maximising light capture for the pinnate-leaved species living under the canopy. Pinnae distributed along a longer axis can reach farther from the plant’s standing point and reduce self-shading in closed habitats. On the other hand, most species with palmate and costapalmate leaves in which the leaflets are arranged radially, are as tall or taller than the cannopy they inhabit (Fig. S2) where light availability is high. However, we found no association between rachis length and shape for pinnate vs. palmate species, thus rejecting **H3** (Fig. 2, Table S8). The lack of association between height over canopy and shape when comparing pinnate vs. palmate-leaved species is consistent with these results, meaning that neither the plant’s position with respect to the canopy nor the rachis length explain leaf shape. The association between rachis length and shape is only significant when comparing pinnate- vs. entire-leaved species, which can be explained by Corner’s rule: large plants have large leaves which necessarily have longer rachises.

### Leaf shape evolution

We found that dissected lineages appeared frequently during the history of palms and from entire-leaved lineages (Fig. 3, S6-7). Additionally, palmate (palmate+costapalmate) lineages appeared at least twice from pinnate lineages, a result consistent with Horn et al. (2009). Using 178 taxa with pinnate or palmate species only, their study concluded that shape is homoplasious and changes between states are frequent. Overall, leaf shape is particularly labile among pinnate-leaved palms and evolutionary changes are frequent between pinnate-entire and pinnate-dissected, involving polymorphism as well.

Based on our ancestral state estimation, the origin of palm leaves is not entirely resolved. We found clear support for an ancestral pinnate leaf, but conflicting results for dissection. On the one hand, entire-leaved states are repeatedly inferred at the root across our models. However, pinnate-entire leaves are almost entirely absent from early diversification of palms, instead dominated by pinnate-dissected leaves (Fig. 4a,b). Palmate and costapalmate leaf shapes appeared between 75-85 Mya, which corresponds to the stem of Coryphoideae. The appearance of palmate shapes roughly coincides with the age of the costapalmate fossil *Sabalites carolinensis* (Santonian, 86.3-83.6 Mya; Berry, 1914), commonly cited as the earliest palm leaf fossil. However, uncertainty as to the stratigraphic assessment of the formation from which *S. carolinensis* was reported calls for the reassessment of the fossil’s age to be Campanian (83.6-72.1 Mya; Greenwood et al., 2022). If that is the case, the oldest costapalmate (here merged with palmate) fossil would postdate the older Coniacian–early Campanian “pinnate-entire” *Phoenicites imperialis* (Dawson) emend. Greenwood et Conran comb. nov. (89.8 to approximately 80 Mya; Greenwood et al., 2022) and be contemporaneous to the mid-Campanian “undivided” *Plicatophyllum* (81-76 Mya; Crabtree 1987; Greenwood et al., 2022). Macrofossils remain rare and provide at best a minimum age for any taxon or morphological character; the absence of older fossils is likely explained by the incomplete nature of the fossil record and its identification. Nevertheless, results on root state estimations should be interpreted with caution since validating them would require a comprehensive taxonomic sampling beyond Arecaceae and our analyses do not consider potential correlations between leaf shape and other morphological characters.

One interesting outcome of our five-state model is the role played by polymorphic lineages. Polymorphism – pinnate-dissected and pinnate-entire – appears as a transitional state between these shapes forming an evolutionary bridge between lineages with non-polymorphic leaf shapes (Fig. 3). Maximum Likelihood estimates of transition parameters also suggest a strong directionality; the highest transition rates being from polymorphic toward pinnate shapes and the second highest was transitions from entire-leaved shapes towards polymorphism (Fig. 4a and S3). As a result, polymorphic lineages are maintained in our ancestral state estimation throughout time despite that only 3% of extant species are polymorphic. Dissected and entire-leaved of pinnate and palmate shapes do not have significantly different speciation rates but the speciation rate for polymorphic lineages is more than twice higher, which also translated into a higher net diversification rate. Thus, according to our best model, the high number of extant pinnate-dissected lineages (1363 species) does not result from its high diversification rates. It could instead be a result of either a high speciation rate of polymorphic lineages with frequent independent transitions to pinnate-dissected leaves or an early success during diversification that facilitated the accumulation of pinnate-dissected lineages throughout the Cenozoic. In either case, the prevalence of pinnate-dissected leaves is an ancient and deeply rooted pattern in the group’s history.

### Leaf shape through time

We found no clear relationship between temperature variations and changes in the relative frequency of leaf shapes, thus we reject **H4** (Fig. 4c). The late Cretaceous and early Eocene were periods of high global temperatures, with ever-wet tropical areas distributed towards high latitudes. Palms thrived during these periods forming the Palmae Province, and becoming ecologically dominant in South America, Africa and India (Pan et al., 2006). During the Eocene, global temperatures decreased at a relatively fast pace (Zachos et al., 2001) and kept declining towards the present, except during the Mid-Miocene climatic optimum. But there is also evidence of aridification across continents over the past 40-50 My alongside temperature variation, leading to the contraction of tropical-like forests towards the equatorial latitudes. The pollen record has revealed patterns of palm extinctions including groups such as Mauritiinae and Eugeissoneae, typical rainforest groups and pinnate-dissected (Bacon et al. 2022, Lim et al. 2021). From our results, this period corresponds to the diversification of entire-leaved lineages, increasing in relative frequency at the expense of pinnate-dissected palms mainly (Figure 4a, b). Global temperature change did not translate into a clear continuous shift in relative leaf shape frequency, but increasing aridity since the Eocene may have promoted the diversification of entire-leaved palms, considering the positive association between precipitation and pinnate-dissected shapes we found in modern palms.

## Conclusion

We explore the drivers of leaf shape evolution in palms by disentangling the associations between shape, climate, and allometric variables related to plant height, and by reconstructing the evolution of shape throughout their evolutionary history. We highlight the importance of considering biotic (intrinsic) and abiotic (extrinsic) factors when studying the evolution of traits. Regarding palms, tall plants are more likely to be dissected and this tendency is mostly driven by pinnate species but not palmate species. High annual precipitation increases the likelihood of having pinnate leaves when controlling for plant height and temperature was not significantly associated with shape. Palms are important representative taxa of tropical forests with more than 2,500 species distributed globally. Exploring how their leaf shape diversity emerged contributes to our understanding of shape and its adaptive potential to future climate conditions.

## Data Availability Statement

All data generated during this study and the coding scripts used to carry out analyses are available as Supplementary Material in the following repository: doi.org/10.6084/m9.figshare.21230453. Chelsa Bioclimate data are available at https://chelsa-climate.org/downloads/ and can be downloaded using the wget command. PalmTraits v1.0 containing morphological traits for all palm species is available from Kissling et al. (2019), Scientific Data. The Global Vegetation Height raster from Simard et al. (2011; Biogeosciences) is available at https://www.sciencebase.gov/catalog/item/50f59de2e4b0114312ab0280. Scripts for data curation are available at https://github.com/mftorres/palm_leaf

## Supplementary Material: Palmleaf_suppmat_20230417.zip

− **palm_leaf_shape_20221201** contains the leaf shape annotations
− **palms_alltraits_curated_20220620 c**ontains the median variables annotated by species
− **SuppMat**

− Figure S1 Correlation coefficients across all variables.
− Figure S2 Species median distributions by leaf shape.
− Figure S3 Paramenters of the best MuSSE model for entire- and dissected-leaved states.
− Figure S4 Paramenters of the best MuSSE model for pinnate- and palmate-leaved states.
− Figure S5 Paramenters of the best MuSSE model for the five-state model.
− Figure S6 Entire- and dissected-leaved ancestral state estimation across palms.
− Figure S7 Pinnate- and palmate-leaved ancestral state estimation across palms.
− Figure S8 Posterior distribution of parameters for the MuSSE model for entire- and dissected-leaved states.
− Figure S9 Posterior distribution of parameters for the MuSSE model for pinnate- and palmate-leaved states.
− Figure S10 Five-trait ancestral state estimation across palms.
− Figure S11 State probability at the root based on the best-fitting model for entire- and dissected-leaved states.
− Figure S12 State probability at the root based on the best-fitting model for pinnate- and palmate-leaved states.
− Figure S13 Map of species distribution centroids for all palms
− Figure S14 Map of global distribution of plant heights for all palm species by leaf shape.
− **Supplementary publication tables**

− Table S1 Number and percentage of species by leaf shape as used in the Generalised Linear Mixed Models.
− Table S2 Palms exhibiting within-species leaf shape variation.
− Table S3 Species not considered in the analyses. Most species in the list are climbing or have climbing and no-climbing habits.
− Table S4 Selected predictive variables used to fit the Bayesian phylogenetic Generalised Linear Mixed Models.
− Table S5 Median (Minimum - Maximum) and range (delta) of species medians by shape and variable for species used in the Generalised Linear Mixed Models.
− Table S6 Median (Minimum - Maximum) and range (delta) of species medians by shape and variable for species used in the five-state ancestral state estimation.
− Table S7 Median (Minimum - Maximum) and range (delta) of species medians by shape and dissection for species used in the two-state ancestral state estimations.
− Table S8 Median, effective sample sizes, and 95% confidence intervals for the estimates in all Generalised Linear Mixed Models.
− Table S9 Entire-leaved versus dissected leaves. Medians and standard deviation of parameter estimates for alternative ML runs on the best fitting model.
− Table S10 Pinnate versus palmate shapes. Medians and standard deviation of parameter estimates for alternative ML runs on the best fitting model.
− **SupplementaryMaterial_20230105 contains supplementary methods and figure legends**

